# Multi-site identification and generalization of clusters of walking behaviors in individuals with chronic stroke and neurotypical controls

**DOI:** 10.1101/2023.05.11.540385

**Authors:** Natalia Sánchez, Nicolas Schweighofer, Sara J. Mulroy, Ryan T. Roemmich, Trisha M. Kesar, Gelsy Torres-Oviedo, Beth E. Fisher, James M. Finley, Carolee J. Winstein

## Abstract

**Background:** Walking patterns in stroke survivors are highly heterogeneous, which poses a challenge in systematizing treatment prescriptions for walking rehabilitation interventions.

**Objective:** We used bilateral spatiotemporal and force data during walking to create a multi-site research sample to: 1) identify clusters of walking behaviors in people post-stroke and neurotypical controls, and 2) determine the generalizability of these walking clusters across different research sites. We hypothesized that participants post-stroke will have different walking impairments resulting in different clusters of walking behaviors, which are also different from control participants.

**Methods:** We gathered data from 81 post-stroke participants across four research sites and collected data from 31 control participants. Using sparse K-means clustering, we identified walking clusters based on 17 spatiotemporal and force variables. We analyzed the biomechanical features within each cluster to characterize cluster-specific walking behaviors. We also assessed the generalizability of the clusters using a leave-one-out approach.

**Results:** We identified four stroke clusters: a fast and asymmetric cluster, a moderate speed and asymmetric cluster, a slow cluster with frontal plane force asymmetries, and a slow and symmetric cluster. We also identified a moderate speed and symmetric gait cluster composed of controls and participants post-stroke. The moderate speed and asymmetric stroke cluster did not generalize across sites.

**Conclusions:** Although post-stroke walking patterns are heterogenous, these patterns can be systematically classified into distinct clusters based on spatiotemporal and force data. Future interventions could target the key features that characterize each cluster to increase the efficacy of interventions to improve mobility in people post-stroke.

## Introduction

Walking patterns differ between individuals to the point that we can recognize people by how they walk ^1^. These individual differences are more marked after a neurologic injury, such as stroke ^2^, due to heterogeneity in stroke lesion type, size, location, and differences in recovery ^3–6^. These individual differences in walking patterns make systematizing treatment prescription for walking rehabilitation interventions a difficult clinical endeavor. The heterogeneity in walking patterns has been acknowledged by Knutsson and Richards as early as 1979, stating that “therapy and training in the hemiparetic patient should preferably be adapted to the disturbance in each individual case” ^4^. In this study ^4^, researchers used paretic electromyography (EMG) and walking kinematics to qualitatively identify three subgroups of abnormal muscle activation during walking post-stroke – early triceps surae activation, decreased activation of paretic musculature, and paretic muscle coactivation. Similarly, Olney and Richards qualitatively identified different subgroups of walking impairments in the spatiotemporal, kinematic, and kinetic domains ^7^. Also, a seminal study by our co-authors* investigated paretic leg kinematics and used hierarchical clustering to identify four clusters of walking behaviors, distinguished by walking speed and paretic knee flexion angles at different points during the gait cycle ^5^. While these studies identified individual differences in paretic leg function during walking, walking patterns post-stroke are defined not only by paretic leg function but also by non-paretic leg function ^8^.

To characterize functional differences between the paretic and non-paretic legs during walking, recent studies have used asymmetry measures in kinematics ^9–14^ or forces ^11,15–18^. These asymmetry measures are compared to asymmetries observed in neurotypical controls to classify individuals post-stroke into subgroups with higher, comparable, or lower asymmetries than control participants ^9–19^. However, individuals might be classified as symmetric in one feature, for example step lengths, while remaining asymmetric in other features, for example joint kinematics ^19^. Therefore, we reasoned that using an approach that allowed simultaneously assessing multiple measures of walking, we can quantitatively identify distinct subgroups of walking behaviors beyond those identified via a single asymmetry measure, which could inform more specific clinical rehabilitation targets.

Identifying subgroups of walking behaviors requires a heterogeneous sample that encompasses the different combinations of gait deviations observed in survivors of stroke. Achieving this heterogeneity necessitates a large sample size. However, studies in walking post-stroke typically collect small, single site samples: for example, a systematic review of 46 studies in walking post-stroke reported sample sizes between 8 and 39 participants ^20^ that lack the heterogeneity of walking behaviors observed post-stroke. Additionally, activity levels, socioeconomic, and ethnic disparities in post-stroke care across geographical locations influence the walking patterns of the samples engaged in research at different sites ^21–25^. Therefore, combining data across different sites increases sample size and heterogeneity of behaviors measured in research studies, which ultimately can improve the generalizability of research findings to the overall post-stroke population.

Here, we obtained measures derived from ground reaction forces (GRFs), which is the simplest data collected across research labs that use instrumented treadmills, to generate a multi-site sample and used a data-driven approach to identify subgroups of walking behaviors in people post-stroke and controls. We used sparse K-means clustering to obtain a subset of features that define walking clusters, and determined whether different levels of function and impairment distinguish these walking clusters, such that they are indicative of different walking subgroups. We determined whether the observed walking clusters are generalizable across research sites. We hypothesized that participants post-stroke will have different walking features resulting in different clusters of walking behaviors, which are also different from control participants ^5,14,26^. Our results could provide the basis for designing and testing targeted interventions aimed at improving walking quality in people post-stroke.

## Methods

### Data Curation

The lack of standardized protocols for collecting, processing, and analyzing walking data can limit researchers’ ability to combine walking data ^27^. However, bilateral ground reaction forces (GRFs) measured using instrumented treadmills are commonly collected across research sites. These GRFs result from muscles generating forces at each segment which then is applied to the ground during walking ^28,29^, providing insight into how each extremity contributes to the main objectives of walking, defined as shock absorption, stance stability, and forward progression ^29^. Additionally, these GRFs can be used to derive spatiotemporal walking metrics such as step lengths, step times, and speed ^30^. Thus, we gathered GRF data from individuals with chronic hemiparetic stroke walking at their self-selected speed. We gathered data from: Rancho Los Amigos (Rancho, N=7 ^31^), collected from an overground force plate at 2500 Hz (Kistler Instrument Corp., Amherst, NY) for the paretic extremity only. University of Southern California (USC2018 N=22 ^9^ and USC2021 N=23 ^12^, the first author reviewed all data to ensure no duplicates between studies): participants walked for three minutes on a Bertec instrumented treadmill (Columbus, OH, USA) that measured ground reaction forces at 1000 Hz. Kennedy Krieger Institute/Johns Hopkins University School of Medicine (JHU, N=10 ^19^): participants walked for five minutes on a Motek instrumented treadmill (Amsterdam, NL) that recorded forces at 1000 Hz. Emory University (Em, N=9 ^32^): participants walked for one minute on a Bertec instrumented treadmill (Columbus, OH, USA) that measured ground reaction forces at 1000 Hz. University of Pittsburgh (Pitt, N=21^33–35)^ participants walked between 150 and 320 strides on a Bertec instrumented treadmill (Columbus, OH, USA) that measured ground reaction forces at 1000 Hz (Fig. 1A). We received GRF data for each gait cycle, normalized to 100 points per gait cycle, filtered with a fourth order lowpass Butterworth filter with a 15 Hz cut-off frequency for JHU and with a 20 Hz cutoff for Pitt. Data from Emory were shared as raw data.

**Figure 1.**
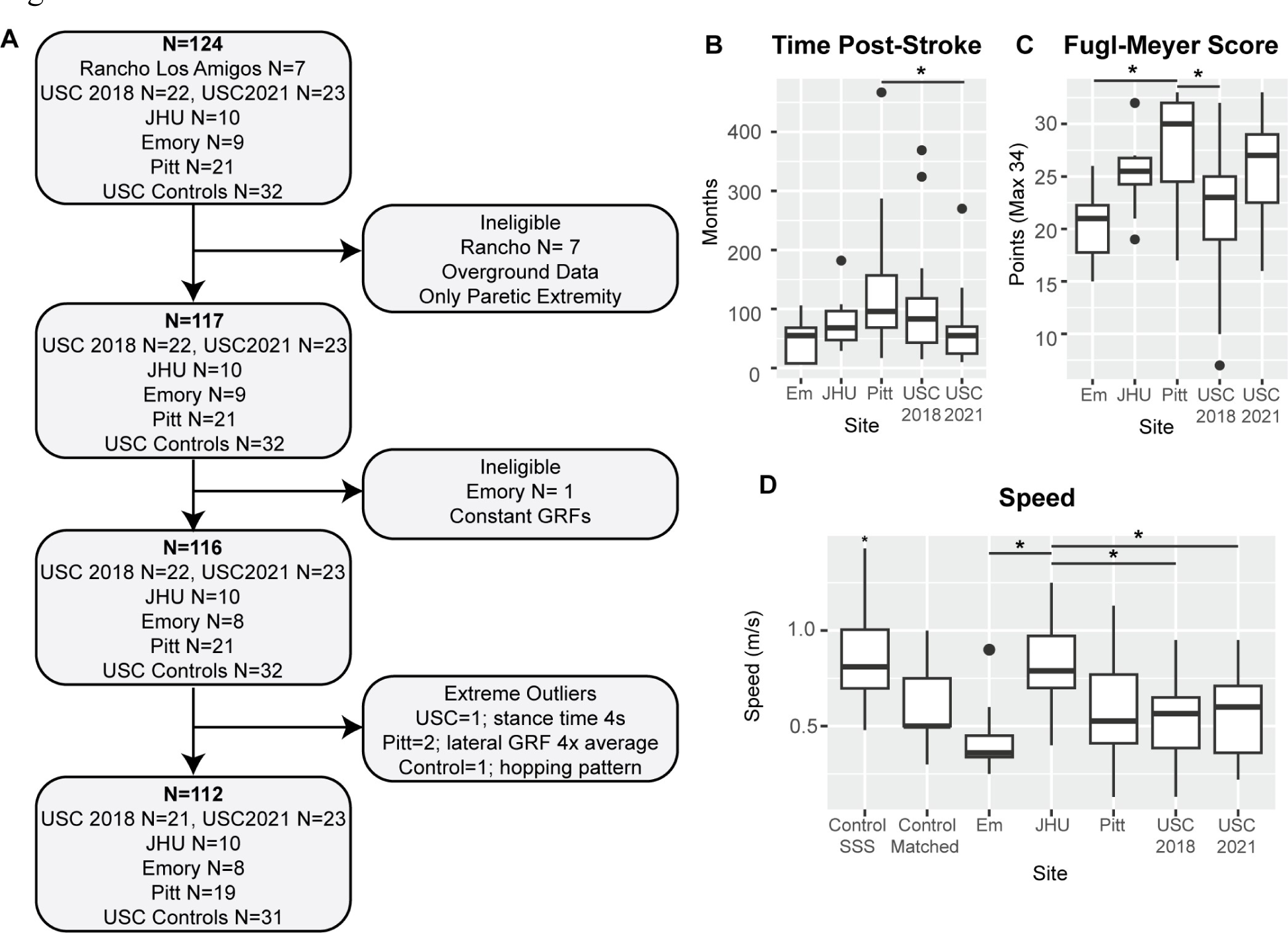
Data consolidation and quality assessments and demographics for the final sample. A) Pipeline for participant inclusion. N indicates the sample size at each point for the total sample (bold) and the sample from each research site. The final sample size was N=112 participants. B) Time post-stroke onset in months across all research sites. Participants from Pitt were more chronic than those from USC2021 (p=0.033). C) Post-stroke participant lower extremity Fugl-Meyer score. The maximum score is 34 points. Participants from Pitt were less impaired than those from Em and USC. D) Treadmill self-selected walking speed for all participants. Control self-selected speed was significantly higher than the average speed in the stroke group (p<0.001). Control participants also walked at speed matched to that of a stroke participant of the same age and sex (Control Matched). Participants from JHU walked at a significantly greater speed than those from Em and both samples from USC (p<0.050).

We filtered data from Em and USC using a 20 Hz cut-off low-pass zero-lag digital Butterworth filter.

We also collected data for 32 age and gender-matched neurotypical control participants using a Bertec instrumented treadmill (Columbus, OH, USA). We used the control data in the definition of the different walking clusters, to determine whether individuals post-stroke could be comparable to neurotypical adults walking at the reduced speed of people post-stroke, such that controls and participants post-stroke would belong to the same clusters. Control participants were age and sex matched to the participants in the USC2021 study ^12^.

Participants post-stroke held onto a front handrail during walking in all studies except USC2021 ^12^ and Pitt ^33–35^. Control participants did not hold onto a handrail. The respective IRB approved all studies, and all participants provided written informed consent before testing. Data collection for neurotypical control participants was approved by the USC IRB number HS-19- 00430. Data gathering was approved by the USC IRB number HS-19-00075. IRB-approved Data Use Agreements (DUA) were established between USC and each research institution. Data from Rancho were excluded as they were collected overground; all DUAs and data sharing manuals were first implemented with Rancho, which was vital in setting up this project. We accumulated GRF data for 92 participants post-stroke (Fig. 1A). We compiled information about participant age, sex, time post-stroke in months, paresis, mass, treadmill walking speed, and lower extremity Fugl-Meyer score ^36^ (out of 34 points for the motor scale) (Table 1).

**Table 1:** Participant Demographics.

|  | **Stroke** | **Control** |
| --- | --- | --- |
| **N** | 81 | 31 |
| **Sex** | 28F/53M<sup>+</sup> | 19F/12M |
| **Age** | 58.9 ± 10.5 [32 – 77] years | 63.3 ± 13.6 [24 – 81] |
| **Mass** | 85 ± 18.6* [47 – 131] kg | 73 ± 15.7 [46 – 110] |
| **Speed** | 0.58 ± 0.24 [ 0.13 – 1.25] m/s | SS: 0.84 ± 0.18* [0.48 – 1.43] m/s<br>Matched: 0.60 ± 0.25 [0.3 – 1.0] m/s |
| **Paresis** | 42R/39L |  |
| **Time post-stroke** | 90 ± 83 [6 – 467] months |  |
| **Fugl-Meyer Score** | 24.6 ± 5.6 [7 – 33] |  |

Descriptive statistics are presented as average ± standard deviation with the range in brackets.
F: Female
M: Males
L: Left
R: Right
SS: Self-selected
\*Significant differences between participants post-stroke and controls (p<0.050)
<sup>+</sup>p<0.050 significant difference in the frequency of males compared to females in the study sample

### Data processing

From the GRF data collected across labs, we used custom code and derived 17 walking variables common across all labs. Variable definitions are presented in Table 2. We obtained averages across strides for all variables for each participant. Data processing was done using custom code written in Matlab 2021b (The MathWorks, Natick, MA).

**Table 2:** Group-level averages for gait variables in participants post-stroke and control participants walking at matched speeds.

| Variable | Definition | Stroke |  | Control |  |
| --- | --- | --- | --- | --- | --- |
| | | Paretic Mean ± SD | Non-Paretic Mean ± SD | Non-dominant Mean ± SD | Dominant Mean ± SD |
| **Speed** | Treadmill speed | 0.58 ± 0.24 | | 0.60 ± 0.18 | |
| **Step Length (m)** | Distance between limbs at leading limb initial contact | 0.37 ± 0.10 | 0.36 ± 0.11 | 0.39 ± 0.09 | 0.39 ± 0.10 |
| **Stance Time (s)** | Time between initial contact and lift-off on the same side | 0.97 ± 0.25<sup>*,+</sup> | 1.00 ± 0.28<sup>+</sup> | 0.86 ± 0.23 | 0.87 ± 0.23 |
| **Swing Time (s)** | Time as the time between lift-off to initial contact on the same side | 0.47 ± 0.09<sup>*</sup> | 0.43 ± 0.07 | 0.45 ± 0.06 | 0.45 ± 0.06 |
| Double Support Time (s) | Time from contralateral initial contact to ipsilateral foot-off | 0.29 ± 0.13<sup>+</sup> | 0.24 ± 0.19 | 0.20 ± 0.08 | 0.21 ± 0.09 |
| **Peak Medial GRF (N/kg)** | medially directed force during the contralateral toe-off<sup>28</sup> | 0.93 ± 0.22<sup>*</sup> | 0.82 ± 0.20 | 0.85 ± 0.28 | 0.82 ± 0.26 |
| **Peak Lateral GRF (N/kg)** | Laterally directed GRF during loading response | -0.24 ± 0.21<sup>*</sup> | -0.39 ± 0.24<sup>+</sup> | -0.20 ± 0.18 | -0.21 ± 0.17 |
| **Peak Propulsive GRF (N/kg)** | Force during ipsilateral push-off | 0.67 ± 0.34<sup>*,+</sup> | 1.03 ± 0.46<sup>+</sup> | 0.87 ± 0.34 | 0.87 ± 0.34 |
| **Peak Braking GRF (N/kg)** | Force during weight acceptance/weight transfer | -0.86 ± 0.44<sup>*</sup> | -0.75 ± 0.44 | -0.80 ± 0.33 | -0.84 ± 0.31 |
| **Vertical GRF (N/kg)** | Force due to gravity | 10.18 ± 1.05 | 10.15 ± 0.99 | 10.2 ± 2.50 | 10.7 ± 2.68 |
| % Gait Cycle of Peak Propulsive GRF | Relative to paretic limb initial contact | 47 ± 12% | 19 ± 22% | 57 ± 9% | 11 ± 12% |
| % Gait Cycle of Peak Braking GRF | Relative to paretic limb initial contact | 16 ± 16% | 52 ± 17% | 16 ± 3% | 66 ± 4% |

t-tests corrected for multiple comparisons using the Benjamini and Hochberg FDR<sup>60</sup>.
\*Significant differences between the paretic and non-paretic extremity of post-stroke participants
<sup>+</sup>Significant differences between participants post-stroke and controls
Variables in bold were included in sparse clustering analyses.

For data collected in neurotypical adults, leg dominance was defined as the leg they would use to kick a ball, which was the right leg for all participants. We compared the non-dominant leg of control participants to the paretic leg and the dominant leg to the non-paretic leg.

Stride-by-stride data in a subset of participants post-stroke and controls are available for download from the Stroke Initiative for Gait Data Evaluation (STRIDE) database ^37^ hosted by the Archive of Data on Disability to Enable Policy and Research (ADDEP).

### Statistical analyses

Statistical analyses were done in RStudio, R version 4.1.2. Since multiple researchers collected data, identifying potential acquisition issues and quality assessment was done post-hoc. To remove noisy data and outliers, we quantified the mean and standard deviation of all variables in our sample, including both participants post-stroke and controls and removed participants with datapoints outside of the mean ± 3*standard deviation range (Supplementary Table 1). The final sample comprised 81 participants post-stroke and 31 controls for 112 participants (Fig. 1A). For all analyses comparing participants post-stroke with neurotypical controls, we used data with neurotypical control participants walking at a matched speed to participants post-stroke.

Given that many walking variables are correlated due to the inherent coordination found in the walking pattern, the clusters found in the data can be accounted for by a small subset of features. Thus, we used sparse K-means clustering analyses ^38^, which uses a Lasso penalty ^39^ to derive a sparse subset of features with non-zero predictors. We z-scored all variables before analyses, and set K=5 clusters in agreement with prior work identifying four subgroups of participants post-stroke ^5^, and an additional control group. We verified that K=5 provided clusters that maximized the between clusters distance via the Krzanowski and Lai index^40^. Using the sparcl package in R, we chose the tuning parameter for the Lasso penalty ^39^, which determines the number of non-zero predictors to use in our analyses that maximizes between cluster variance and minimizes within cluster variance. We ran 100 permutations in a search space between 1 and the square root of the number of candidate variables included in the sparse clustering analyses, i.e., sqrt(17) and set the number of random starts to 100 to avoid finding a local minimum.

We assessed stability of the clusters identified in this study via the Jaccard similarity index ^41–43^. We resampled the 112 participants with replacement via bootstrap to obtain 10,000 new samples, and identified K=5 clusters for each bootstrap iteration. We then measured the proportion of observations consistently assigned to the same cluster over each iteration; this proportion is the Jaccard index. A Jaccard below 0.6 indicates that the clusters are unstable, a Jaccard between 0.6 and 0.75 shows moderately stable clusters, between 0.75 and 0.85 stable clusters, and a Jaccard above 0.85 indicates highly stable clusters.

We used linear models with *cluster* as a categorical fixed effect to determine whether there were significant differences across clusters in participants’ biomechanical features, as well as participant demographics and Fugl-Meyer scores. We performed multiple comparisons using Tukey-Kramer adjusted critical values. We used the results from these multiple comparisons to characterize the cluster-specific walking behaviors, by identifying which features were significantly different across clusters. Within each cluster, we used t-tests with false discovery rate (FDR) corrected p-values for all spatiotemporal and force variables, to compare paretic values relative to non-paretic values to determine asymmetries in each cluster.

We assessed generalizability of each research site by determining the number of participants from each site that were included in each cluster. To assess whether the observed clusters can be generalized, we performed a leave-one-out sparse K-means clustering approach, leaving out data from each site simultaneously (Em, Pitt, JHU, USC2018, USC2021). We mapped the clusters obtained in each iteration of the leave-one-out approach to the clusters using the full dataset. We interpret cluster generalizability based on its Jaccard ^43^ and the similarity of the descriptive walking and impairment measures within each cluster.

## Results

Our final sample was composed of 112 participants (Fig. 1, Table 1 and 2, Supplementary Fig. 1). At a group level, participants post-stroke showed significant differences in paretic swing (p=0.007) and stance times (p=0.005), medial and lateral ground reaction forces (p<0.001), braking (p=0.032), and propulsion (p<0.001) compared to the non-paretic extremity. Chronicity (p=0.024), impairment measured via FM score (p=0.001), and walking speed (p<0.001) differed across sites (Fig. 1B-D).

Sparse K-means clustering identified 8/17 features with non-zero weights from which five distinct clusters could be obtained (Fig. 2, Supplementary Figs. 2 – 5). The Krzanowski and Lai index (Supplementary Fig. 2) was maximal for K=5 clusters. Analyses of the clusters obtained showed different walking behaviors from those observed at a group level. The variables that maximized between cluster variance and minimized within cluster variance, as shown by the non-zero weights returned by the Lasso analyses, were paretic and non-paretic stance times, non-paretic propulsion, speed, paretic step length, paretic and non-paretic braking, and non-paretic step length (Fig. 2). The data for the paretic extremity was combined with the non-dominant extremity in controls, and the data for the non-paretic extremity was combined with the dominant extremity in controls. We describe the obtained clusters next:

**Figure 2.**
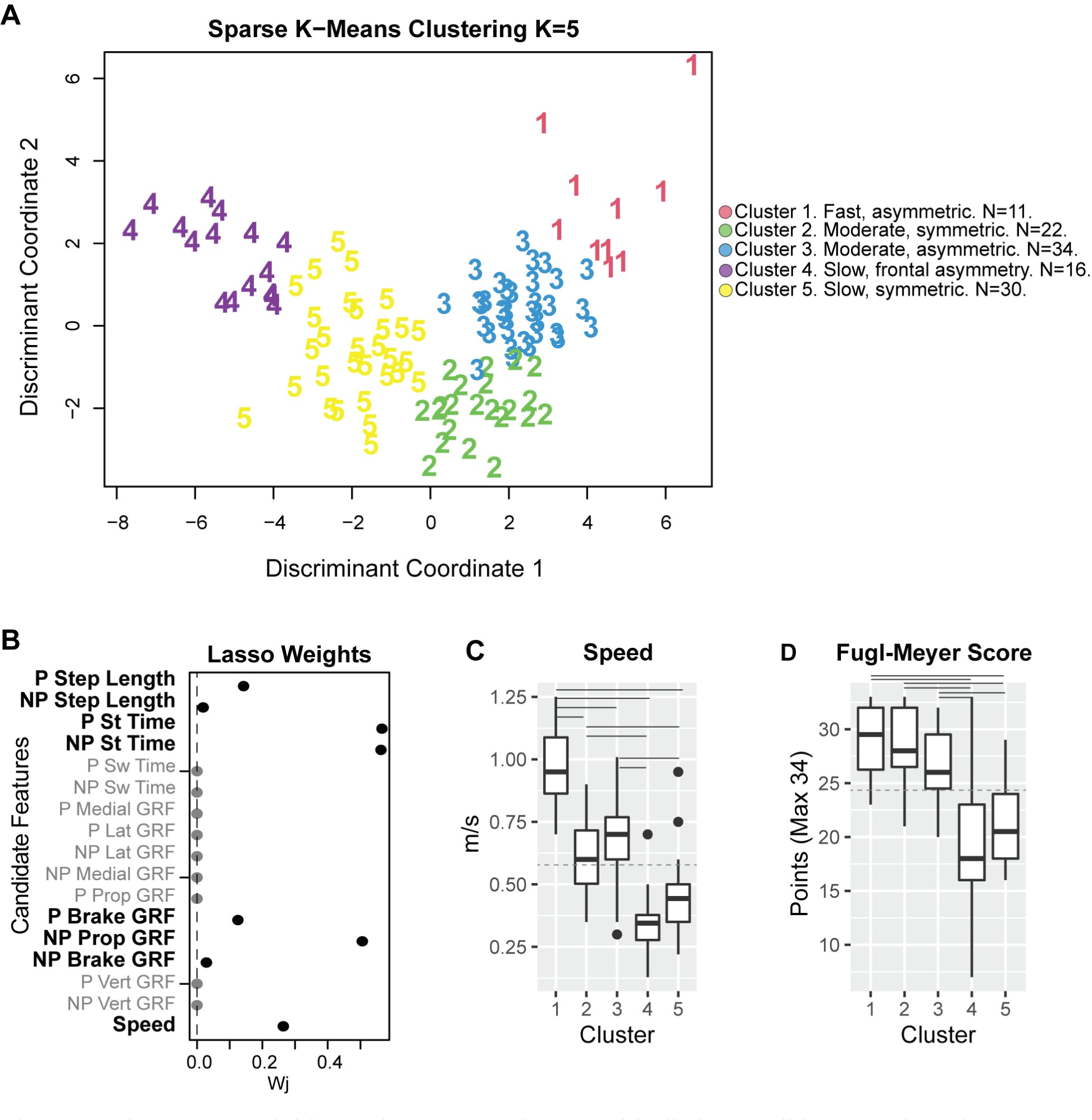
The sparse variables make up K=5 clusters with distinct walking speeds and impairments. A) Weights for each of the 17 candidate features for clustering. 8/17 features had non-zero weights and were used in clustering analyses. B) Individual observations within each cluster, plotted in discriminant component space, colored by cluster. Cluster 1 comprised 10 participants; four from JHU, three from Pitt, two from USC2018, and one from USC2021. Cluster 2, 22 participants; 11 controls, four from Pitt, one from USC2018, and six from USC2021. Cluster 3, 34 participants; 11 control participants, one from Em, four from JHU, four from Pitt, seven from USC2018, and seven from USC2021. Cluster 4 had 16 participants; three controls, three from Em, five from Pitt, three from USC2018, and three from USC2021. Cluster 5 had 30 participants; six controls, four from Em, two from JHU, three from Pitt, seven from USC2018, and seven from USC2021. C) Participant speed (for participants post-stroke and controls) and impairment (for stroke participants) were measured via FM score across the different clusters. Only speed was used in the cluster definition. Solid horizontal lines indicate post-hoc significant differences between clusters (p<0.050). The dashed horizontal line indicates group-level average speed across all participants and average Fugl-Meyer (FM) scores for all stroke participants compared with the cluster-level speed and FM. P: Paretic NP: Non-Paretic. St: Stance. Sw: Swing. Lat: Lateral. Prop: Propulsion. GRF: Ground reaction force.

### Cluster 1 – Fast speed and asymmetric stroke cluster

N=10 stroke participants. Composed of four participants from JHU, three from Pitt, two from USC2018 and one from USC2021. Participants had mild impairment (FM 29±3, Fig. 2D).

#### Walking characteristics

Participants in the fast and asymmetric cluster had the fastest speed (0.97±0.17 m/s, p<0.001, Fig. 2C, Fig. 3), and were classified as community ambulators ^6,44^. Step lengths were 0.49±0.05 and 0.51±0.07 m for the paretic and non-paretic extremities, the longest of all post-stroke participants. Participants had the largest braking and propulsive GRFs bilaterally (p<0.001), and showed marked asymmetries: non-paretic propulsion was 72% greater than paretic propulsion (p<0.001), and non-paretic stance (p=0.015) and paretic swing times (p=0.017) were longer than the contralateral extremity.

**Figure 3.**
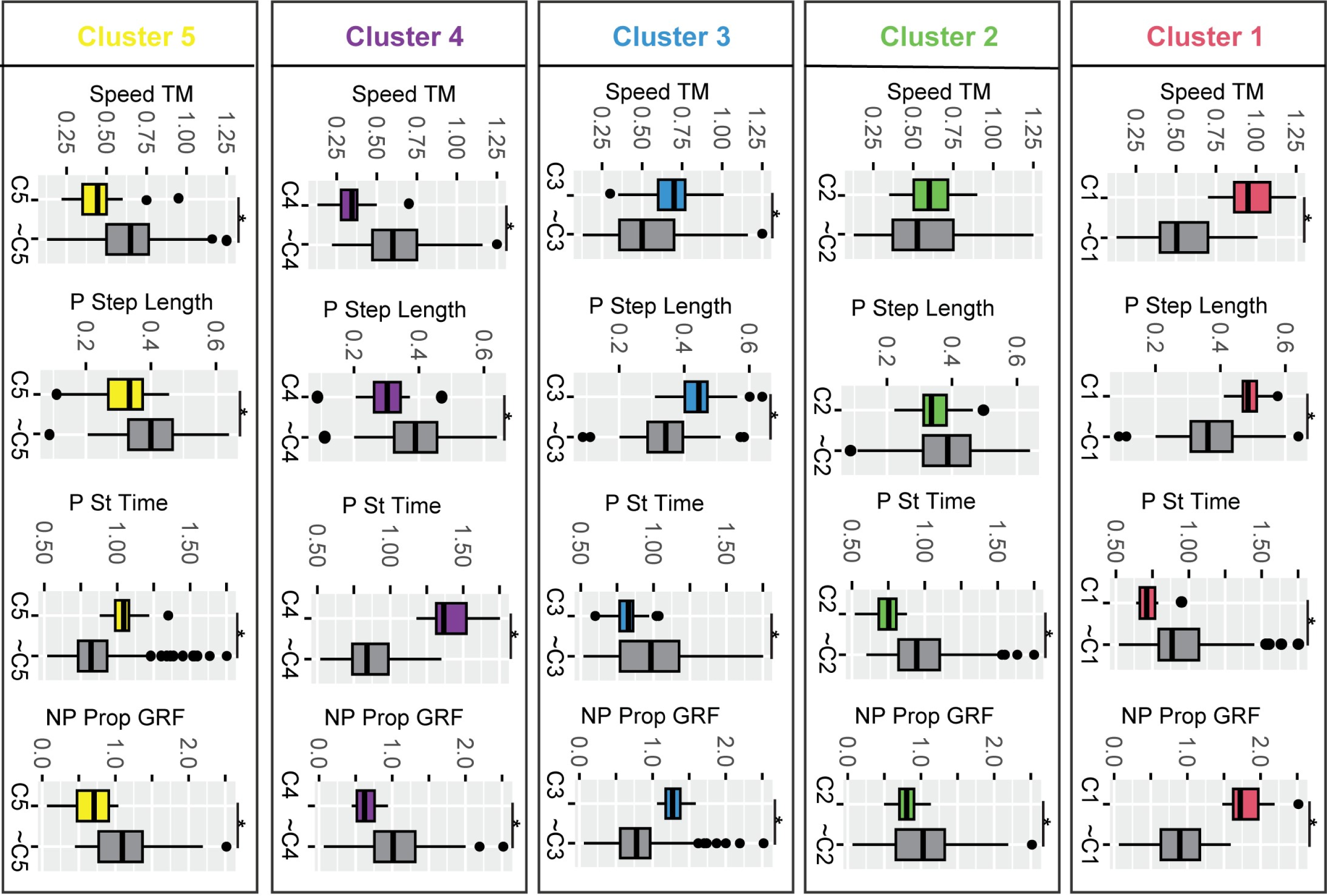
Gait features for participants post-stroke and controls within each cluster compared to all other participants outside each cluster for a subset of sparse variables used in k-means clustering. C1, … C5 indicates participants included in the respective cluster number, and **∼**C1, … ∼C5 indicates all participants not included in the respective cluster. Stance and swing times are expressed as a percentage of stride duration to account for differences in walking speed. Cluster 1: participants post-stroke with a fast-walking speed and asymmetric propulsion; Cluster 2: participants post-stroke and controls with moderate speed, short stance times, low propulsion, and symmetric steps. Cluster 3: participants post-stroke and controls with moderate speed, short stance times, and asymmetric forces; Cluster 4: participants post-stroke with a slow speed and frontal plane force asymmetries; and Cluster 5: post-stroke participants who walked slowly and symmetrically, with short swing times. Color conventions as in Figure 3. * FDR corrected p<0.010 for all variables indicated as significant. Abbreviations as in Figure 2.

#### Cluster stability and generalizability

The Jaccard index for the fast and asymmetric cluster was 0.72 (Fig. 4), and during the leave-one-out validation varied between 0.63 and 0.75. Leaving out the data from JHU, which was the sample with the highest walking speed, had the greatest effect on cluster stability and reduced the Jaccard to 0.63. Given the absence of the JHU participants, many participants changed cluster membership and the number of participants in the fast and asymmetric cluster increased to 32, and the average speed decreased to 0.78m/s as the speed requirement for this cluster became less stringent. Despite the change in membership, the general characteristics of the fast and asymmetric cluster remained consistent, thus, the cluster is generalizable.

**Figure 4.**
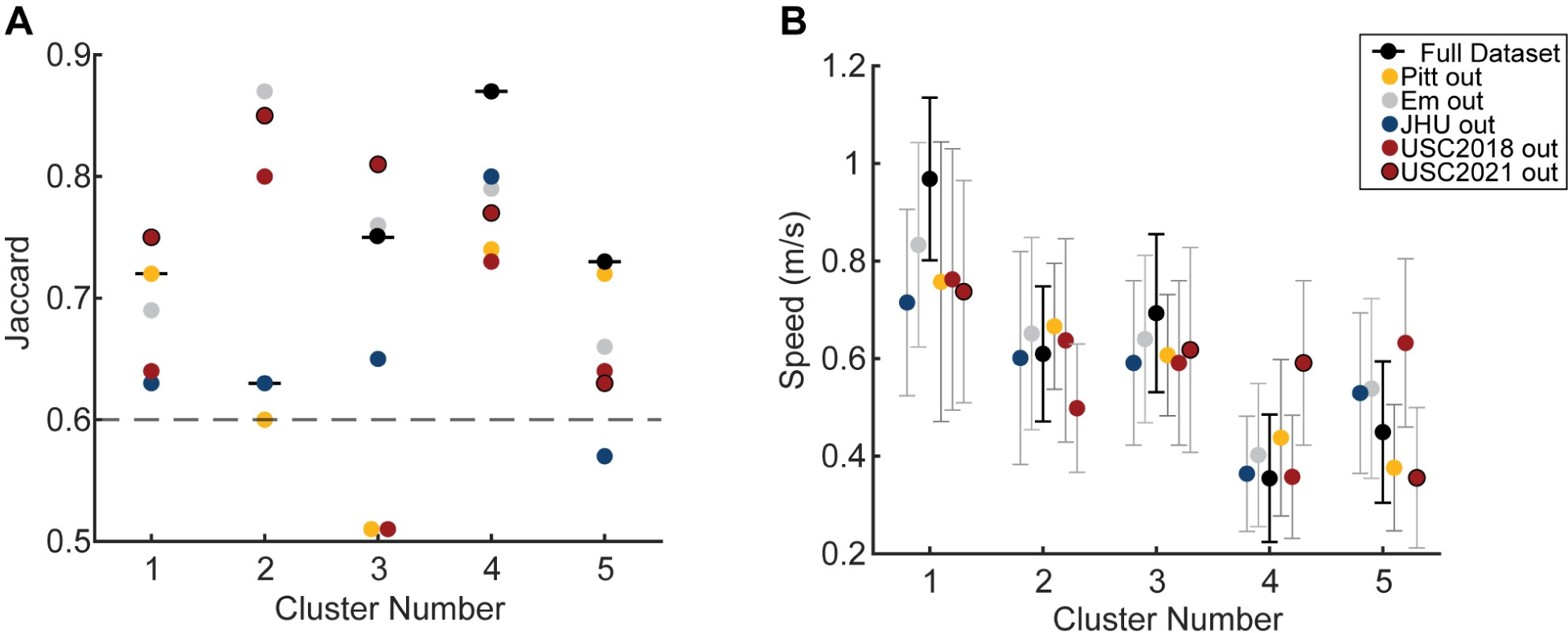
Stability and generalizability of clusters assessed via leave one out approach. A) Jaccard stability index for each cluster for the entire dataset (horizontal black lines) and leaving out each sample. Colors indicate the sample left out to assess the Jaccard. The dashed gray line is the line above which clusters are considered stable. The Jaccard Index is the proportion of observations consistently assigned to the same cluster over the bootstrap iterations. A Jaccard below 0.6 indicates that the clusters are unstable, a Jaccard between 0.6 and 0.75 shows moderately stable clusters, 0.75 and 0.85 stable clusters, and a Jaccard above 0.85 indicates highly stable clusters. B) Average and standard deviation of the speed for each cluster for the entire dataset in black and the leave one out approach with colors indicating the sample left out from analyses.

### Cluster 2– Moderate speed, symmetric and short stance times in controls and stroke

N=22 composed of 11 control and 11 post-stroke participants. Participants post-stroke were four from Pitt, one from USC2018 and six from USC2021. Participants post-stroke in the moderate speed, symmetric cluster had mild impairment (FM 29±4, Fig. 2D).

#### Walking characteristics

Participants in the moderate speed, symmetric cluster walked at 0.61±0.14 m/s (greater than the slow clusters, p<0.005 and less than the fast cluster, p<0.001), classified as least limited community ambulators ^6,44^. Step lengths were 0.35±0.06 m for both the paretic and non-paretic extremities. Additional analyses also showed that stance times were shorter in the moderate speed, symmetric cluster compared to all other clusters (Fig. 3). Post-stroke participants in the moderate speed, symmetric cluster were symmetric across all variables measured, making them comparable to controls walking at this slow speed.

#### Cluster stability and generalizability

The Jaccard index for the moderate speed, symmetric cluster was 0.63 and varied between 0.6 and 0.87 during validation, bordering on stable to highly stable. Leaving out the data from Em, JHU, USC2018, and USC2021 made the moderate speed, symmetric cluster exclusively a control cluster with Jaccard indices of 0.87, 0.63, 0.80, and 0.85. Leaving out the data from Pitt reduced the Jaccard to 0.6 and reduced the number of control participants in the moderate speed, symmetric cluster to eight, with one participant from Em, four from JHU and four from USC. Given that this cluster is primarily a control cluster during validation and that the Jaccard remains in the moderately stable to the highly stable range, the moderate speed, symmetric cluster is generalizable and may indicate changes in the walking pattern that are adopted when people walk at a slower speed.

### Cluster 3 – Moderate speed, asymmetric cluster of participants post-stroke and controls

N=34. 11 controls and 23 participants post-stroke from all sites. Post-stroke participants had mild impairment (FM 27±3, Fig. 2D).

#### Walking characteristics

Participants in the moderate, asymmetric cluster walked at 0.69±0.16 m/s, classified as least limited community ambulators ^6,44^ (faster than Cluster 4 and Cluster 5, p<0.005 and slower than Cluster 1, p<0.001). This was the only cluster that showed cluster-level step length asymmetry, with longer paretic steps (p=0.032).

Stance times were a shorter percentage of the stride sample compared to all other clusters (Fig. 3). Forces were highly asymmetric both for participants post-stroke and control participants in this cluster. The non-paretic peak lateral GRF was 62% greater than the paretic lateral GRF (p<0.001), non-paretic propulsion was 62% greater than paretic propulsion (p<0.001), and paretic braking was 28% greater than non-paretic braking (p=0.001).

#### Cluster stability and generalizability

The Jaccard index for the moderate, asymmetric cluster using the entire dataset was 0.75 and during the leave-one-out validation varied between 0.51 and 0.81, from unstable to stable. Leaving out the data from Pitt or USC2018 reduced Jaccard indices to 0.51 (Fig. 4A). Leaving out the data from Em, JHU, USC2018, or USC2021 made the moderate, asymmetric cluster a primarily control cluster composed of three participants post-stroke from Pitt and nine controls. In contrast, leaving out the data from Pitt made the moderate, asymmetric cluster a primarily stroke cluster with three control participants and 13 participants post-stroke. Given the change in cluster from a control cluster to a stroke cluster during validation, this cluster is not generalizable across sites.

### Cluster 4 – Slow speed and frontal plane force asymmetries

N=16 primarily stroke participants, three from Em, five from Pitt, three from USC2018, two from USC2021 with moderate to mild impairment (FM 20±8, Fig. 2D). This cluster included three control participants who walked with a frontal plane waddling pattern to account for the slower speeds to match participants post-stroke.

#### Walking characteristics

Participants in the slow cluster with frontal plane asymmetry walked at 0.35±0.13 m/s (slower than the fast and the two moderate speed clusters, p<0.001), classified as unlimited household ambulators ^6,44^. Step lengths were 0.29±0.08 and 0.31±0.09m for the paretic and non-paretic extremities, respectively. Participants within the slow cluster with frontal plane asymmetry had stance times that were a longer percentage of the gait cycle compared to all other participants (p<0.001), non-paretic stance was markedly longer than paretic stance (p<0.001), and paretic swing time was also longer than non-paretic swing time (p=0.038). The paretic medial GRF was 25% greater than the non-paretic counterpart (p=0.013), and a paretic lateral GRF that was half of its non-paretic counterpart (p=0.005, Fig. 3). Non-paretic propulsion was 61% greater than paretic propulsion (p=0.006). Participants in the slow cluster with frontal plane asymmetry generated non-paretic propulsion at around 31% of the gait cycle, significantly later than all other clusters (p<0.0001), and had significantly longer double support times than all other clusters (p<0.001).

#### Cluster stability and generalizability

The Jaccard index for the slow cluster with frontal plane asymmetry using the entire dataset was 0.87, and during the leave-one-out validation varied between 0.73 and 0.80, indicating a stable cluster. Given the high stability with the entire dataset and maintained stability during the leave-one-out approach, this cluster is generalizable across sites.

### Cluster 5 – Slow speed, symmetric

N=30, primarily stroke cluster of participants from all sites with moderate impairment (FM 21±4, Fig. 2D).

#### Walking characteristics

Participants in the slow speed symmetric cluster walked at 0.45±0.13 m/s (slower than the fast and moderate speed clusters, p<0.001), classified as the most limited community ambulators ^6,44^. Step lengths were 0.31±0.08 and 0.31±0.09m for the paretic and non-paretic extremities, respectively. The percent of the gait cycle in the swing phase was significantly shorter for participants in the slow speed, symmetric cluster. Participants within the slow speed symmetric cluster were symmetric except with regards to propulsion, which was 30% greater in the non-paretic extremity (p=0.005), and the medial GRF, which was greater in the paretic extremity (p=0.003).

#### Cluster stability and generalizability

The Jaccard index in the slow speed, symmetric cluster using the entire dataset was 0.73 and during the leave-one-out validation varied between 0.57 and 0.72, from unstable to moderately stable. Leaving out the data from JHU, which contributed two participants to this cluster, had the greatest effect on cluster stability, increasing the Jaccard to 0.57 and walking speed to 0.60 m/s. Leaving out the USC2018 data decreased the Jaccard to 0.64 and increased walking speed to 0.63 m/s. Despite the lower Jaccard, the characteristics of the slow speed symmetric cluster, mainly low propulsion and short swing times remained consistent, making this cluster generalizable.

## Discussion

We identified five clusters of walking behaviors in a combined sample of people post-stroke and controls. Contrary to our hypothesis, we did not identify a cluster composed exclusively of control individuals: we identified a cluster with an equal number of control participants and participants post-stroke, the latter of whom seemed to have reduced walking impairment. We also identified clusters with mostly participants post-stroke and a few controls, which points to the fact that slow walking speeds can lead to aberrant gait patterns even in healthy controls ^45^. The clusters obtained in our study point at different levels of function and impairment defining each cluster, which correspond to different walking subgroups post-stroke. We assessed the generalizability of clusters and observed that 4/5 clusters were generalizable across research sites. These clusters were: Cluster 1: a cluster of post-stroke participants with fast walking speed and asymmetric propulsion; Cluster 2: a cluster of controls and stroke participants walking with moderate speed and symmetric steps, with apparent impairments due to reduced speed, such as short stance times, low propulsion; Cluster 4: a cluster of participants post-stroke walking with a slow speed and asymmetric medial and lateral ground reaction forces; and Cluster 5: a cluster of post-stroke participants who walked with a slow speed, short swing times and slow but symmetric steps. It may be possible to develop more personalized intervention targets by considering the cluster to which a given patient is assigned. For example, a post-stroke participant in the fast cluster can benefit from an intervention to increase paretic propulsion ^17,46–48^; a post-stroke participant in the moderate symmetric cluster can benefit from an intervention to increase speed ^13,49–51;^ a post-stroke participant in the slow, frontal asymmetry cluster can benefit from interventions to reduce asymmetries in medial and lateral GRFs ^52^; and a post-stroke participant in the slow symmetric cluster can benefit from increasing speed ^53^, swing times, and step lengths. The effects of these targeted interventions are still unknown, and future work will determine whether addressing cluster specific impairments will lead to a shift towards a control cluster, or a shift towards a different stroke cluster. Taken together, our results provide a preliminary cross-sectional analyses to estimate which subject-specific walking-related variables could be targeted in interventions to improve mobility and promote neuroplasticity in the long term ^54^.

Our findings complement those of our co-authors who showed speed to be the greatest determinant in allocating participants to specific clusters ^5^, and previous work using speed alone to classify participants to ambulation categories ^6^. Similar to Mulroy, 2003, we found one fast cluster of participants post-stroke, one moderate speed cluster and two slow clusters. A recent study used paretic and non-paretic kinematic data in 36 stroke survivors and identified six distinct walking clusters ^55^ based on range of motion for the paretic and non-paretic side. Like our results, Kim et al. observed different types of impairment associated with the different clusters. A potential limitation of these two studies is that they comprise single-site datasets, which might be biased by geographical affordances. We complement the clusters described by Mulroy by providing insights into both paretic and non-paretic spatiotemporal characteristics and forces generated by each limb. One of our slow clusters had asymmetric mediolateral forces (Cluster 4), which corresponds to the slow extended cluster in Mulroy 2003: this cluster in Mulroy 2003 showed knee hyperextension in mid-stance, which limits pre-swing and swing knee flexion and toe clearance, leading to frontal plane compensations and asymmetric frontal plane forces, as observed in our study. Our other slow cluster, Cluster 5 also corresponds to the very slow velocity and excessive knee flexion cluster from Mulroy 2003. Thus, comparison of our clusters to those previously reported show consistency of subgroups of walking patterns, and provide additional insights into non-paretic function in these subgroups.

The variables that had the largest effect on the between cluster variance, were paretic and non-paretic stance times, non-paretic propulsion, speed, paretic step length, paretic and non-paretic braking, and non-paretic step length. It is worth noting that none of these variables on their own are sufficient to identify the clusters that we observed, similar to what was concluded by Mulroy, 2003. For example, speed ranges overlapped for participants in Clusters 2 and 3 and for participants in Clusters 4 and 5, yet different stance times and forces were observed between clusters despite overlapping speeds, indicating that speed alone was not the only factor driving the between cluster differences. Thus, here we show that while speed influences many gait features, speed alone is not enough to classify post-stroke individuals in more specific subgroups of impairment.

Cluster 3, the moderate speed and asymmetric cluster was not stable or generalizable, consistent with the fact that the common impairments of stroke participants, and how they compare to control participants within Cluster 3 was less clear. For example, participants in Cluster 3 had highly asymmetric forces between the paretic and non-paretic extremity whereas control participants within the cluster were not asymmetric. Similarly, at a group level, participants within Cluster 3 were the only ones to show marked step length asymmetry with longer paretic steps. It could be the case that Cluster 3 is composed of participants for whom the measured spatiotemporal and kinetic variables included in analyses does not account for variability in the data. Future work will aim to use joint level kinematic, kinetic or EMG measures to determine whether these more specific measures can detect evident differences in walking patterns in these individuals.

We observed differences in cluster specific speed and FM scores compared to group level averages in these variables. For example, the average FM score for all participants was 24.6 ± 5.6 points, yet as seen in Figure 2D, this average value does not align with any of the cluster-specific averages. We also observed that similar levels of impairment measured via overlapping FM scores between clusters could be associated with vastly different walking speeds: participants in Cluster 1 walked significantly faster than those in Clusters 2 and 3, despite no differences in FM scores. Clinically this might imply that participants in Clusters 2 and 3 have the capacity to walk at faster speeds, given their impairment. Note that these differences in behavior and capacity between clusters could inform clinical practice beyond group level averages.

Interestingly, our findings show that some commonly reported spatiotemporal impairments post-stroke are observed in only a subset of post-stroke participants ^8,28,61,66^. For example, we observed asymmetric stance times only for participants in the fast cluster, and asymmetries in step lengths only in Cluster 3 ^11,61,66,67^, as well as varying degrees of paretic propulsion that were not uniformly associated with walking speed or impairment ^56^. Finally, and surprisingly, we did not observe any asymmetries in the peak vertical GRF among our participants. Thus, our results show in a sample of 81 participants post-stroke that spatiotemporal asymmetries are less pronounced than what has been shown in the seminal literature using smaller samples.

We observed significant differences in participant impairment and function across research sites. Consistent with prior literature, control participants’ self-selected walking speed was faster than post-stroke participants’^7^. Similarly, participants from Pitt had lower impairment compared to Emory and USC2018, while participants from JHU walked at faster speeds than those from Em, and USC. Additional information about activity levels and access to post-stroke care across geographical locations could provide information about the causes of the differences in impairment and function between sites. Given these differences, we assessed generalizability by assessing cluster stability when removing each experimental sample from our dataset. This approach assessed both generalizability of the cluster such that its definition did not depend on the experimental samples, as well as generalizability of the experimental samples such that if a sample needed its own cluster, it would imply that participants in that sample are distinct from all other participants. We did not find a cluster of participants post-stroke from a single research site. Additionally, 4/5 clusters were generalizable across research sites. However, we did find that some research sites did not contribute to specific clusters. For example, participants from JHU were not part of Cluster 4, and participants from Em were not part of the fast cluster, consistent with the significant differences in walking speeds observed between both samples.

These results confirm that some samples may not encompass all different walking behaviors observed after stroke.

## Limitations

There are multiple limitations in our study. First, this study used variables derived from gait analyses to identify clusters indicative of subgroups of walking impairment. The variables used in this study are easy to capture via GRFs and represent global measures of walking that relate to the coordinated action of multiple segments and joints. However, joint-level kinematic and kinetic measures might capture impairments that cannot be inferred from GRF data alone. Future research efforts should aim to establish standards for data collection that allow combining more complex data across sites, such as joint-level kinematics, kinetics and EMG patterns.

Second, we extracted peak force as the feature representing GRF data due to the ease to capture these measures. Future work can assess whether other GRF features, such as impulses, or GRFs during more specific points during the gait cycle provide more insight into the range of walking impairments in people post-stroke. Third, we were also limited in the sample size of control participants which was unbalanced and matched in speed to the USC2021 sample only. The sample size across study sites was also unbalanced, with samples from Em and JHU consisting of ∼10 participants and the samples from USC and Pitt consisting of ∼20 participants, providing different heterogeneity within each sample and different contribution of each site to the overall sample. Stability analyses show however, that changes in stability when removing each of the samples were not just due to sample size: removing the JHU sample which consisted of 10 participants with the highest speeds decreased cluster stability for most clusters, while removing the USC2018 sample did not change cluster stability uniformly. We interpret this to indicate that the characteristics of the samples had a greater influence on cluster stability than did sample size. Future multi-site studies can address these points. Fourth, participants in this study were all in the chronic phase of stroke recovery; future work should assess if these clusters are consistent or change across recovery phases. Fifth, data were collected with participants walking on a treadmill which may induce changes in walking patterns compared to overground walking ^57^.

However, some of our clusters were similar to those reported by Mulroy 2003 ^5^, indicating consistency of clusters on the treadmill compared to overground. In addition, the use of a handrail in some of the experimental protocols could have influenced participant’s walking patterns ^58^. Future work could systematically assess whether participants are assigned to the same clusters measured during treadmill walking as to during overground walking. Finally, we used average metrics within participants, despite different patterns of stride-to-stride variance during post-stroke walking ^13^. Future work in larger samples can include stride-to-stride variance as additional features to characterize post-stroke walking patterns.

## Conclusions

We compiled and curated GRF data across multiple research sites in people post-stroke and controls. Using simple measures derived from GRFs, we identified five clusters of different walking behaviors. Four of these clusters captured walking subgroups that were generalizable across study sites. Our findings provide new information about how to classify the heterogeneity of gait patterns post-stroke. Identifying more specific types of walking impairment and different intervention targets for each subgroup can move the field of neurorehabilitation toward a precision medicine approach ^59^, and improve the effectiveness of rehabilitation interventions.

## Acknowledgments

We want to thank Chang Liu, PhD, Sungwoo Park, PhD, Tara Cornwell, Ryan Novotny, Catherine Yunis, Carly Sombric, PhD, Digna DeKam, PhD, Thu Nguyen, PhD, Justin Liu, Purnima Padmadaban and all trainees who contributed to data collection at all study sites.

This work was funded by:

NIH NCATS grants nos. KL2TR001854 (N Sánchez), R03TR004248 (N Sánchez), and NIH NCMRR grant no. P2CHD065702 (N Sánchez);

NIH NICHD grant no. R01-HD091184 (J.M. Finley),

NIH NINDS grant no. R21 NS120274 (N. Schweighofer) NIH NIA grant no. R21 AG059184 (R. Roemmich)

NIH NINCHD grant nos. R01 HD095975 and R21 HD095138 (T. Kesar)

## Supplementary Materials

**Supplementary Table 1.** Demographics for participants excluded from analyses.

| Group | Speed (m/s) | Mass* (kg) | Age (years) | Sex | FM Score | Affected Side | Months Post-Stroke |
| --- | --- | --- | --- | --- | --- | --- | --- |
| Emory | 0.45 | 71.4 | 74 | F | 26 | L | 24 |
| Pitt | 1.11 | 74.5 | 75 | M | 32 | R | 370 |
| Pitt | 0.53 | 53.9 | 66 | F | 26 | R | 117 |
| USC | 0.13 | 67.0 | 28 | F | 19 | L | 25 |
| Control | 1.05 (matched 0.13) | 51.4 | 28 | F |  |  |  |

\*p<0.05 significant differences for the independent samples t-test comparing the sample of participants excluded vs. the sample of participants included in analyses

**Supplementary Figure 1.**
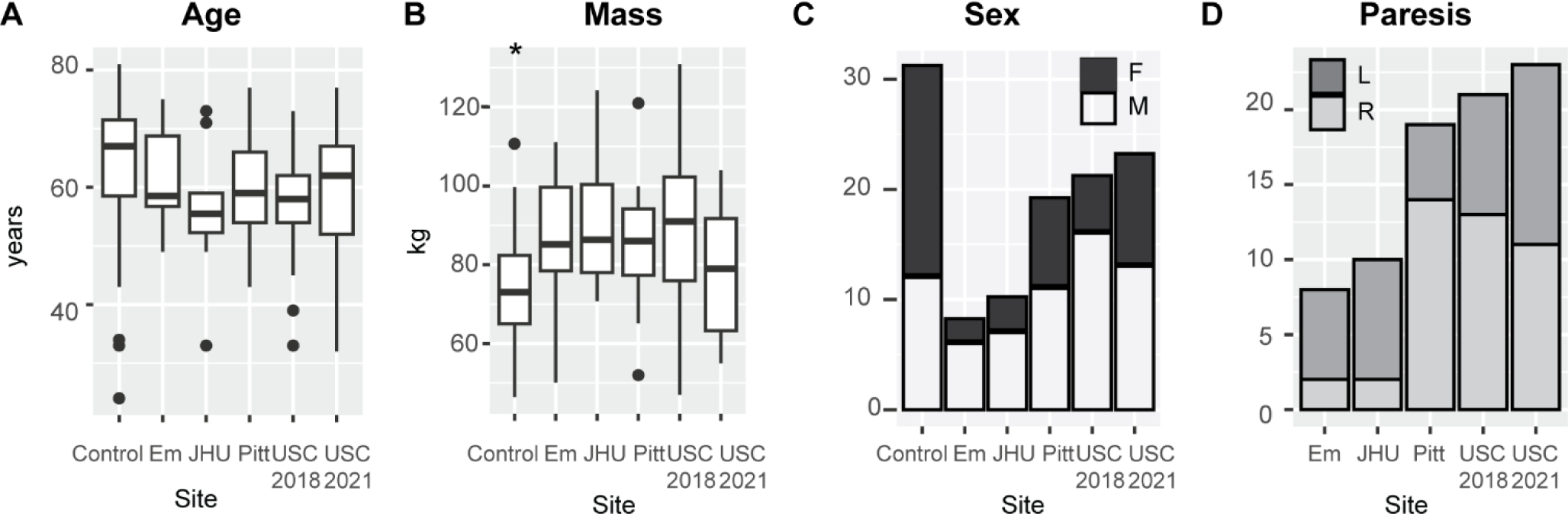
Demographics. A) Participant age in years across all sites tested. B) Participant mass in kilograms. Mass in control participants was significantly lower than in post-stroke participants (p=0.002). C) Self-reported sex across different sites. D) Paretic side for participants post-stroke across different sites.

**Supplementary Figure 2.**
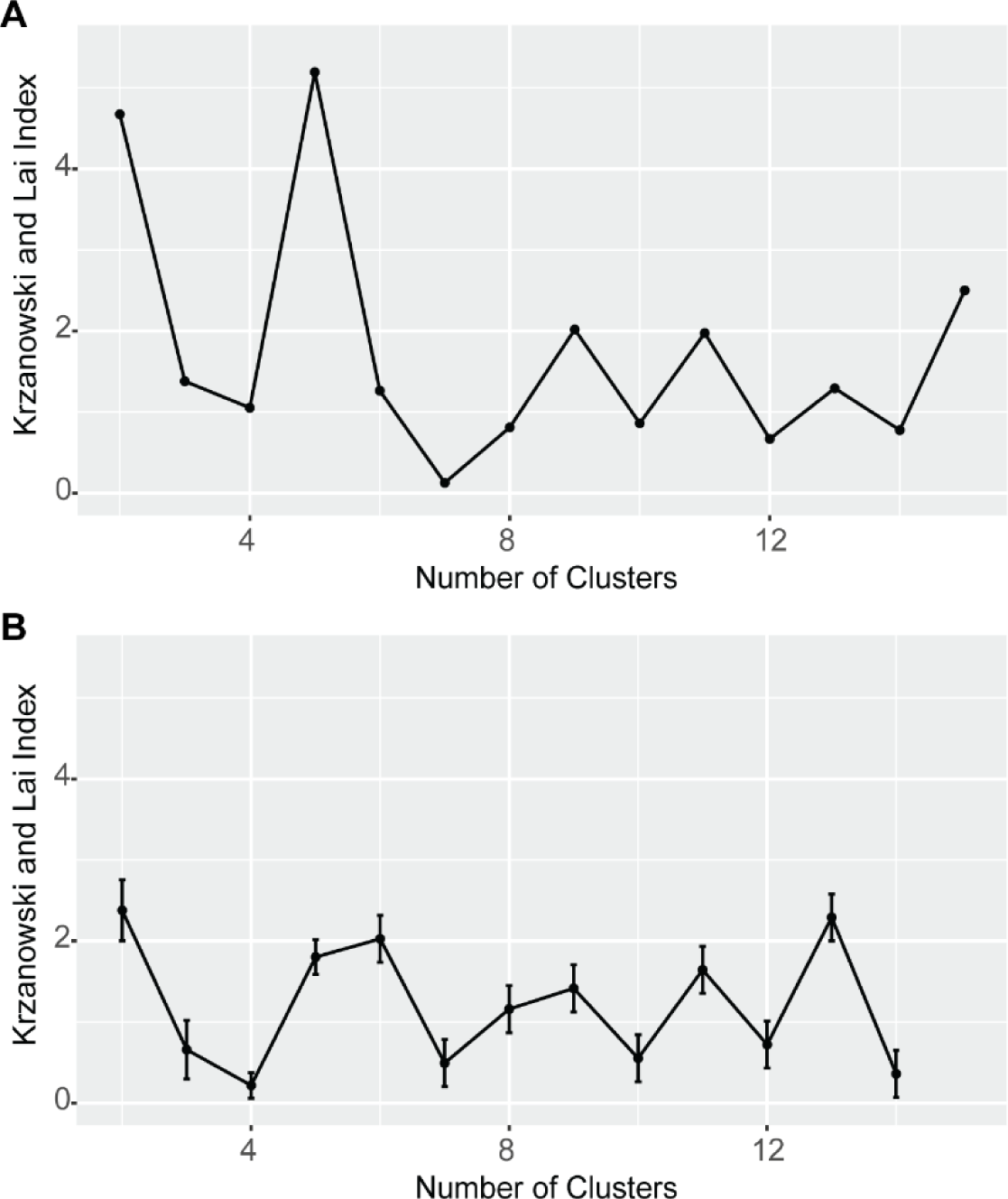
A) Krzanowski and Lai index for D-Index plots for the dataset in our study. The optimal number of clusters is the one with the highest index. B) Mean and 95% confidence intervals for the Krzanowski and Lai index obtained via 1000 bootstrap iterations. Bootstrap analyses indicate overlapping confidence intervals for the Krzanowski and Lai index for 2, 5, 6 and 13 clusters, which are also higher than all other number of clusters. Given previous work identifying 4 stroke clusters plus our group of control participants we maintain K=5 clusters.

**Supplementary Figure 3.**
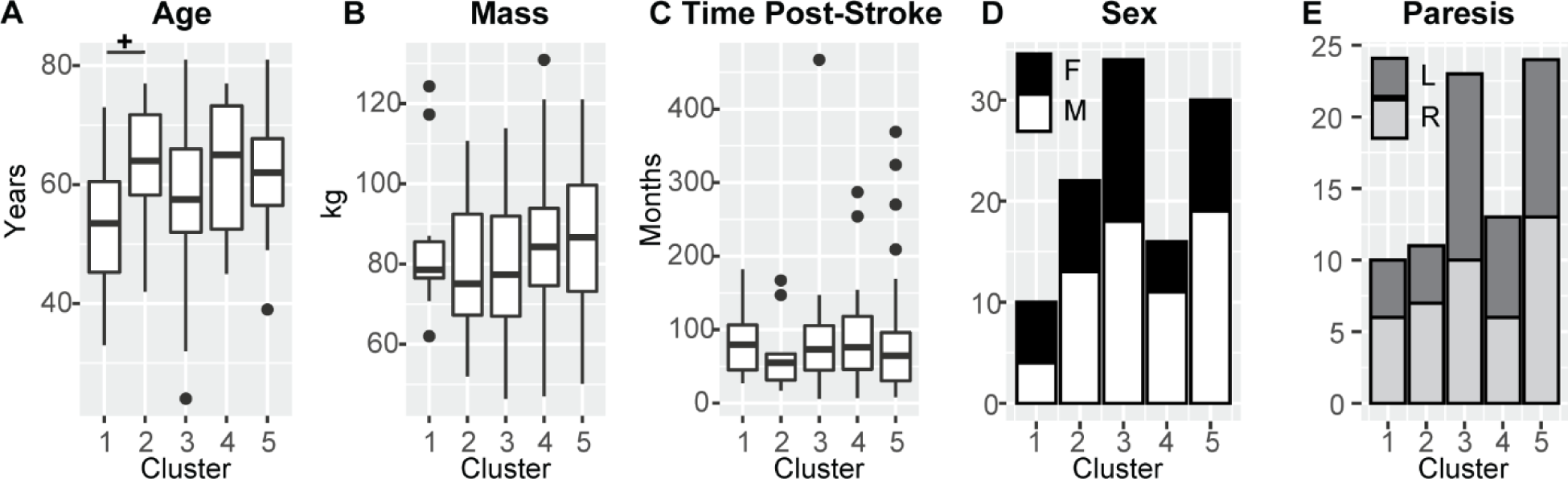
Participant demographics across the different clusters. A) Age in years for participants post-stroke and controls in each cluster. Participants in C1 were marginally younger than C2 (p=0.051). B) Mass in kg for participants post-stroke and controls across clusters. C) Time post-stroke in months for participants post-stroke. No differences were observed across clusters (p=0.925). Horizontal lines indicate post-hoc differences between clusters. D) Sex for participants post-stroke and controls across clusters for all participants. E) Paresis across clusters for participants post-stroke.

**Supplementary Figure 4.**
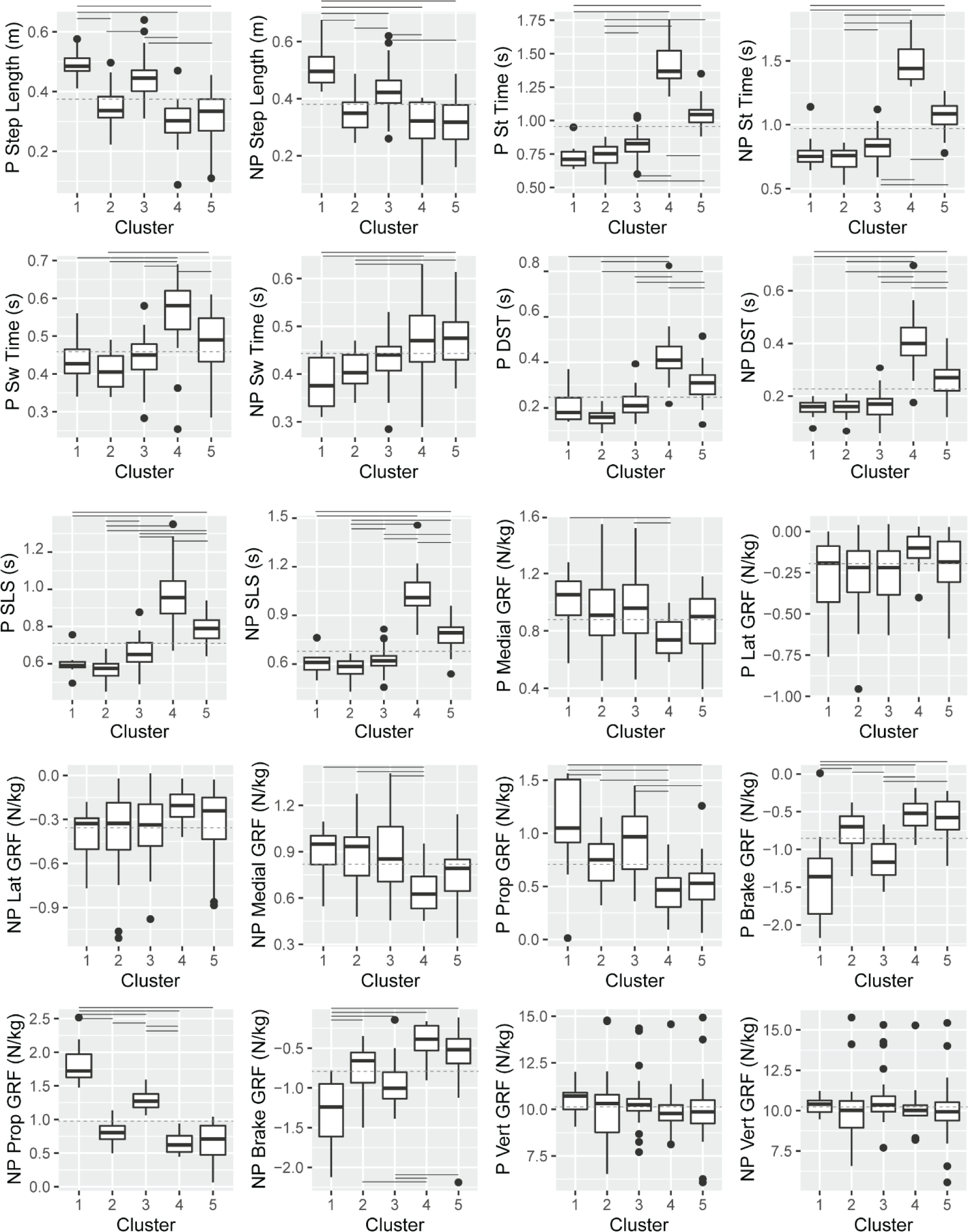
Spatiotemporal and peak forces across the different clusters for participants post-stroke and controls. Post-hoc significant differences between clusters are indicated by the solid horizontal lines (p<0.050). The dashed horizontal line indicates the average value of the variable across all 112 participants to allow comparisons of group-level averages with cluster-level averages. DST refers to double support time, and SLS refers to single limb support time, which was available for all participants except those from Pitt and therefore were not used as candidate variables for clustering. All other variables were used as candidate variables in sparse analyses.

**Supplementary Figure 5.**
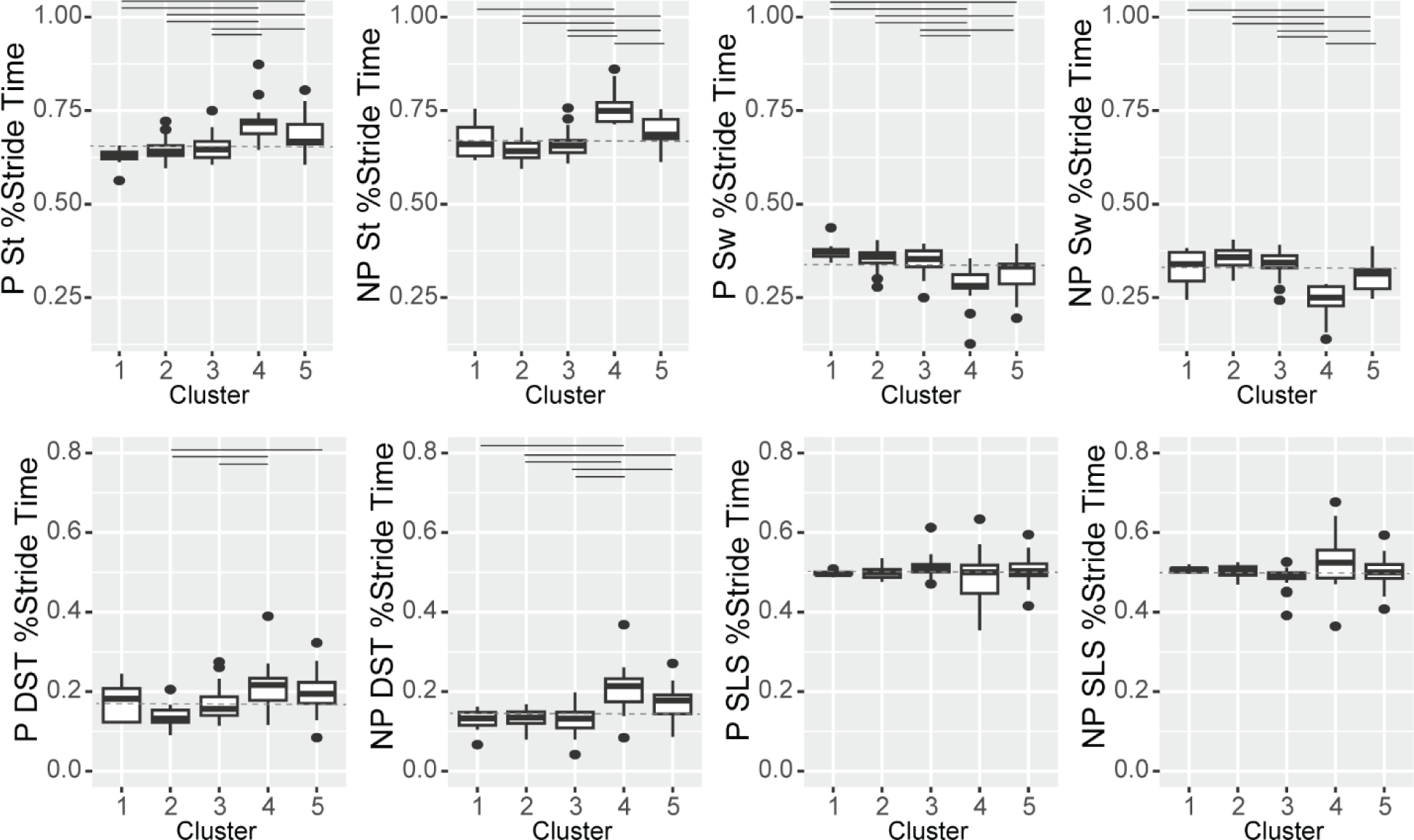
Temporal variables are expressed as the percent duration of the overall stride. Post-hoc significant differences between clusters are indicated by the solid horizontal lines (p<0.050). The dashed horizontal line indicates the average value of the variable across all 112 participants for stance and swing times and 93 participants for double and single support times to allow comparisons of group-level averages with cluster-level averages. DST refers to double support time, and SLS refers to single limb support time, which was available for all participants except those from Pitt.

